# A classification approach to categorize urbanized landscapes in California and trait variation in Maltese starthistle

**DOI:** 10.1101/2025.09.07.674560

**Authors:** Anthony J. Dant, Lily G. Bishop, Katrina M. Dlugosch

**Affiliations:** All University of Arizona, Tucson, AZ, 85721

**Keywords:** Centaurea melitensis, Hierarchical clustering, Landscape ecology, Spatial analysis, Species invasions Urban ecology

## Abstract

There is increasing evidence that the traits of organisms can differ in urban environments, but defining what makes an environment ‘urban’ is difficult. There are many variables associated with increasing human impacts that might be associated with urbanization, and this makes it challenging to identify both how traits vary across diverse urbanized landscapes and the variables that might drive that variation. To better define the mosaic of urban environmental heterogeneity and identify similar environments that are comparable across its complexity, we developed a multivariate landscape classification framework and applied it to classification of land area in the state of California, USA. We then asked if these multivariate Environmental Zones were better able to explain trait variation than univariate urban vs. non-urban dichotomies or gradients, using a case study of the invasive, annual plant Maltese starthistle (*Centaurea melitensis)*, commonly found throughout much of California. We used a hierarchical cluster analysis to group 7,829 government census tracts into Environmental Zones based on a set of 19 independent environmental characteristics, including climate, land cover, pollution, and socio-economic variables. This analysis identified nine major Environmental Zones, which were differentiated based upon complex combinations of variables that did not align with urban vs. natural dichotomies or gradients. Environmental Zones also occurred as mosaics of many zones within cities and differed in their relative abundance between cities, reflecting complex urban landscapes unique to each area. Seeds from seventeen populations of *Maltese starthistle* were collected from six of the Environmental Zones, including two more heavily urbanized zones, and four more natural/agricultural. Seeds were grown in greenhouse conditions, and eight vegetative traits were measured. No trait differed significantly between qualitatively urban versus non-urban sites, or across a quantitative gradient of impervious surface area, but four traits differed according to Environmental Zones (length of longest leaf, SLA, root diameter, and the number of flowerheads). Traits differed between the more urbanized zones, as well as among the more natural zones. Our results reveal that more complex multivariate classifications of the urban mosaic can identify similar, comparable environments across complex landscapes and better explain trait variation in organisms navigating urbanized environments.

## INTRODUCTION

There is increasing recognition that humans have altered landscapes in a wide variety of ways, and that anthropogenic modifications to the environment are increasing (Grimm et al., 2008; Johnson & Munshi-South, 2017; Szulkin et al., 2020). While humans have been creating permanent settlements for as long as 10,000 years (Elmqvist et al., 2013) the manipulation of the natural environment through large-scale agricultural operations and artificial housing structures emerged as recently as 8,000 years ago (Lev-Yadun et al., 2000; Kareiva et al., 2007). Modern urban structures such as extensive impervious surfaces, large scale housing, and mechanical industry, emerged during the late 18th century in Europe and are now increasingly common globally (Elmqvist et al., 2013). Due to the scale at which these human modifications alter landscapes, there is growing scientific interest in how such novel urban environments are affecting the organisms that inhabit these spaces, both non-human and human alike (McPhearson et al., 2016; Rivkin et al., 2019; Szulkin et al., 2020). As a result, there is a growing need for informative methods of quantifying the mosaic of environments created by the interaction between human modifications and the natural landscape (Ramalho & Hobbs, 2012; Winchell et al., 2022; Murray-Stoker et al., 2025). Here we use a classification approach to categorize an environmental mosaic and apply this information to explain trait divergence in an invasive plant that occupies a wide range of urbanized to natural environments.

At first glance, urban environments seem to be simple to define as human population centers distinct from natural areas, offering a straightforward dichotomy for comparisons between urban and natural environments (Sukopp, 1998; Grimm et al., 2008; Niemelä, 2011). Urban environments are envisioned to force abrupt changes on natural processes, with (potentially) easily quantifiable effects on ecology and evolution relative to natural habitats (Pickett et al., 1997; Alberti et al., 2020; Miles et al., 2020). Indeed, in a review of methodology used in 181 studies of urban ecology and evolution, Szulkin and colleagues (2020) found that 75% of studies utilized such qualitative descriptions to define urban areas. Investigating this traditional dichotomy has proven fruitful in uncovering evidence that ecological and evolutionary processes can be different in urban environments (Buczkowski & Richmond, 2012; Marzluff, 2017; Putman & Tippie, 2020; Hou et al., 2023). However, it has become clear that ‘urban’ environments are highly heterogeneous spaces. To understand the impacts of those environments on organisms, we must develop a better understanding of that heterogeneity (Rivkin et al., 2019; Verrelli et al., 2022).

The traditional idea that urban environments are distinct entities has shifted into a more nuanced concept, the urban-to-natural gradient (McDonnell & Hahs, 2008; Ramalho & Hobbs, 2012). The gradient concept describes urban environments as being the epicenter of artificial structures and impervious surfaces, which are then assumed to decline essentially linearly into more natural environments, usually with urban followed by suburban, agricultural, rural, and finally natural areas. The urban-to-natural gradient views urban environments as the center of disturbances that permeate into subsequent environments, based on geographical location (Larondelle & Haase, 2013; McPhearson et al., 2016; Aronson et al., 2017). The appeal of this univariate gradient approach is that it is relatively tractable with these assumptions. Studies of phenotypic changes in the form of morphology, behavior, and physiology have been analyzed through this urban-to-natural framework (Urban et al. 2006; Giraudeau et al. 2014; Ariori et al. 2017; Salmón et al. 2018; Santangelo et al. 2020).

However, problems have arisen from using the gradient concept to classify environments, because it can fail to address the complexities found within each category (McDonnell & Hahs, 2008; Ramalho & Hobbs, 2012; Moll et al., 2019). For example, pollutants such as nitrogen oxide species (a common byproduct of vehicular traffic) within a city may not be equally distributed across all urban areas, thus making some parts of a city less, or potentially even more, habitable for particular species (Sanderfoot & Holloway, 2017; Johnson et al., 2024). Anthropogenic influences also have the potential to interact with each other and their climatic context, forming local microclimates shaped by multiple environmental variables, such as changes in land use and climate, creating a mosaic of environments throughout urban sprawl (Čeplová et al., 2017; Pickett et al., 2017; Peterson et al., 2020).

It is increasingly recognized that a consequence of simplifying the categorization of urban landscapes is that even studies that have identified phenotypes associated with urban environments have then had mixed success in identifying what specific environmental variables lead to these differences (Szulkin et al., 2020; Verrelli et al., 2022). For example, urban birds are particularly well-studied and many species of birds have experienced morphological changes that lead to increased fitness in urban environments, yet few studies have determined the environmental influences underlying these changes (Evans et al. 2009; Marzluff 2017; Matuoka et al. 2020). Additionally, studies of plants have found trait differences between urban and natural areas, e.g. in phenology and biomass (Yakub & Tiffin, 2017; Dubois & Cheptou, 2017; Fukano et al., 2023; Sexton et al., 2023), but the specific environmental factors that have led to these changes remain unclear. Finally, studies across multiple urban areas have looked for evidence of repeated adaptations to similar urban environments, but with mixed results, raising the question of which environments are most comparable (Trentanovi et al. 2013; Diamond et al. 2018; Santangelo et al. 2022). These issues highlight the need to identify which features of urban environments contribute to urban phenotypes.

Finally, socio-economic and socio-cultural elements have been historically excluded from urban ecological and evolutionary studies (Schell et al., 2020). Variability in socio-economic variables across urban environments has been linked to changes in resource availability, habitat modification, and patterns of selection on traits (Munshi-South, 2012; Alberti et al., 2017; Peterson et al., 2020; Verrelli et al., 2022). For example, in arid regions, more affluent areas are associated with greater water availability through their use of irrigation (Diamant et al. 2025). Thus, these variables can influence the environment experienced by organisms (Wood and Esaian 2020, Des Roches et al. 2021, Johnson et al., 2024) but are rarely included in environmental classification models within urban studies.

For these reasons, researchers in urban biology have begun to urge a more detailed multivariate classification of urban environments, so that studies can more accurately identify the environmental heterogeneity found within cities and surrounding areas (Rivkin et al. 2019; Winchell et al. 2022; Leandro et al. 2025). Recent reviews also encourage quantification of urban heterogeneity via spatially acquired data, due to its potential to more accurately explain ecological and evolutionary factors on a landscape scale (Szulkin et al, 2020; Fusco et al. 2021; Murray-Stoker et al. 2025).

To address this need, here we develop a multivariate land classification framework for California, USA, and use this framework to identify trait differentiation in the invasive, annual plant Maltese starthistle (*Centaurea melitensis* L., Asteraceae). We build upon a land classification approach developed through the Urban Nature Research Center at the Natural History Museum of Los Angeles County in Los Angeles, California, USA (Li et al., 2019). This method groups geographical units using a hierarchical cluster analysis, based on quantifying multivariate differences among those units. This approach was originally developed to characterize the distribution of biodiversity across the city of Los Angeles, CA. Here we leverage a similar approach to classify heterogeneity of the entire landscape of urban, rural, and natural areas across the state of California into a set of Environmental Zones, based on an expanded set of variables that include socioeconomic factors. We then apply the Environmental Zone classification to a to test for phenotypic differences across populations of Maltese starthistle. We evaluate whether the new classification framework is more informative for understanding trait variation than existing binary or gradient definitions of urban environments. Our goal is to offer a framework for better understanding the mosaic of variables that result in ecological and evolutionary changes across urban environments.

## METHODS

### Geographic area and units of analysis

California is an ideal region for the development of urban classification approaches because it contains many cities of varying sizes and substantial publicly-available spatial data. The state includes six cities with populations over 500,000 (U.S. Census Bureau 2021), extensive housing developments that directly border natural areas, and a wide diversity of climatic, environmental, and socio-economic conditions (Berg et al., 2024). The State of California consolidates spatial data into accessible public sources, including three databases used in this study: the WorldClim v2.1 database (https://www.worldclim.org), the California Environmental Protection Agency (CALEPA; https://calepa.ca.gov), and the Multi-Resolution Land Characteristics Consortium (MRLC; https://www.mrlc.gov). Within California, we chose the state census tract to be the unit of analysis for this study. Census tracts are the smallest scale at which California spatial vector data are commonly stored. California currently includes 8,057 census tracts, based on the 2020 census (U.S. Census Bureau 2021). We elected to include all census tracts in the state in our analysis, despite our specific interest in urbanization, for several reasons. Foremost, a central goal of our study is to classify human-impacted environments broadly, without a priori assumptions about which areas are sufficiently urbanized to merit study. In addition, classifying areas across the spectrum of human impact and wildland allows us to compare these areas to one other. Finally, analyses at the state scale are practical because data are uniformly available at this scale, and outputs of our study can then be used by the state government for management purposes across the state.

### Environmental variables

Environmental variables available for each census tract were selected across four variable types: land cover, climate, pollution, and socio-economic attributes. Potential variables included 49 variables accessible from the CALEPA (accessed on April 2025), the Multi-Resolution Land Characteristics (MRLC) Consortium (https://www.mrlc.gov, accessed on April 2025), and the WorldClim v2.1 database (https://www.worldclim.org, accessed on April 2025). These variables were then filtered for those that were relevant to non-human terrestrial organisms, those without substantial missing data, and those without strong positive or negative correlations (> |0.7|) making them redundant with other retained variables (see Appendix S1 for all variable selection decisions). This resulted in a total of 19 variables for this study (Table 1). All pollution and socioeconomic variables were stored in pre-existing polygons constructed by the CALEPA (CalEnviroScreen 4.0, 2021).

**TABLE 1.** Environmental variables used in the hierarchical cluster analysis.

|  |  |
| --- | --- |
| Land cover variables<sup>a</sup><br>● Impervious Surface<br>● Tree Cover<br>● Land classification<br>○ Shrubland<br>○ Grassland<br>○ Cropland and Pastureland<br>○ Woody and Herbaceous Wetland | Pollution variables<sup>c</sup><br>● Particulate Matter 2.5<br>● Pesticide Usage<br>● Vehicular Traffic<br>● Cleanup Sites<br>● Groundwater Threats<br>● Impaired Water Bodies<br>● Toxic Releases from Facilities<br>● Diesel Particular Matter |
| Climate variables<sup>b</sup><br>● Maximum Temperature of the Warmest Month<br>● Minimum Temperature of the Coldest Month<br>● Average Annual Precipitation | Socio-economic variables<sup>d</sup><br>● Poverty Index<br>● Asthma Rate |

<sup>a</sup>Source for all land cover variables: United States National Land Cover Database
<sup>b</sup>Source for all climate variables: WorldClim Database
<sup>c</sup>Source for all pollution and socio-economic variables: 2019 CalEnviroScreen 4.0 Report (CALEPA)

Land cover data included several resources available through the MRLC. Impervious surface cover (including concrete, pavement, buildings, and other constructed materials) was obtained through the U.S. Geological Survey (USGS) 2024 Annual NLCD Collection 1 Science Products. Tree cover was obtained through USGS and U.S. Forest Service 2024 National Land Cover Database Tree Canopy Cover Products. All other land cover variables were obtained through the USGS Annual Land Cover database. The amount of each land cover type within each census tract was calculated using the Tabulate Area function in ArcGIS Pro (v3.03). These values were then converted into percentages of the total pixels found within each polygon.

Climatic variables were collected from the WorldClim v2.1 Dataset (Fick and Hijmans 2017) as raster data at 30 seconds resolution (∼1 km2). The average of each variable over the census tract was then calculated using the Zonal Statistic function in ArcGIS Pro. However, some census tracts were too small for this function. If a climatic value was missing from a census tract, the average of all surrounding census tracts was calculated and given to any census tracts which were missing data.

Finally, census tracts were excluded from the study if any variables were not available, with the exception of climatic variables interpolated as described above, resulting in a total of 7,829 out of 8,057 tracts used in this analysis. A PCA was constructed to visualize each of these variables in the dataset, using factoextra (Kassambara & Mundt, 2020) in R (v4.5.0 2025) via RStudio version 4.1.3 (RStudio Team 2023).

### Hierarchical cluster analysis

A hierarchical cluster analysis of similarity among census tracts was conducted using the dendextend (v1.14.0; Galili 2015) and tidyverse (v1.3.0; Wickham et al. 2019) packages in R. Hierarchical clustering treats each unit, in this case a census tract, as a single cluster, which is then merged with other clusters based on their similarities (or distances), until a nested hierarchical structure is formed (Johnson 1967). Here we used Euclidean distances, after standardizing variables to z-scores using the scale function in R (Lele 1993). The sf package (v1.0-12; Pebesma 2018) was used to upload census tract shapefiles in the form of polygons into the analysis with default settings. Large multivariate distances between clusters were identified as long branches in the analysis output dendrogram, and these were designated as unique ‘Environmental Zones’ for this region. We examined the composition of these Environmental Zones by computing the average value of contributing environmental variables within each zone. All 19 environmental variables were log-transformed as log(x+1) prior to calculating means. Minimum Temperature of the Coldest Month was shifted by 12.28 (adding this constant to all values) before transformation to ensure all values were positive. A forest plot of the means of all 19 log-transformed variables was used to visualize the contribution of each variable to each Environmental Zone, relative to each variable’s overall mean. We examined the distribution of zones by computing their representation in different areas of the state, and their representation at different distances to the nearest urban center. Distance to an urban center was calculated as the distance between the centroid of the census tract and its distance (km) to the nearest urban center, defined as a continuous area greater than or equal to 2 km^2^ composed of equal to or greater than 80% impervious surface (see Appendix S2: Supporting Methods - Distance to Urban Center). Linear regressions were used to test for relationships between the percentage of census tracts belonging to an Environmental Zone and the midpoint of 30km bins of distance to the nearest urban center. Finally, we compared the representation of Environmental Zones among the three largest cities in California using a Pearson’s chi-squared test to compare the representation of Environmental Zones in census tracts among Los Angeles County, San Diego County, and the San Francisco Bay Area, including size counties as recognized by the Association of Bay Area Governments including: San Francisco, Alameda, Contra Costa, San Mateo, Marin, and Santa Clara counties.

### Maltese starthistle study system and collections

Maltese starthistle is an invasive annual forb which was introduced to the United States most likely about 150 years ago, as a contaminant of agricultural seeds (Washington State Noxious Weed Control Board 2023; California Invasive Plant Council 2023). Several features make it especially suitable for studies of responses to local urban environments. First, Maltese starthistle currently inhabits a wide range of California habitats, including urban areas of Los Angeles, San Diego, and San Jose. Second, it is an annual that reproduces only by seed, with no clonal reproduction (California Invasive Plant Council 2023), and so it has experienced many generations in its current habitats. Third, it appears to propagate primarily through self-pollinated seeds (California Invasive Plant Council 2023), which likely limits its dispersal across environments, and allows realistic propagation of genotypes in the greenhouse through selfing.

Seeds of Maltese starthistle were collected from across California in the summer of 2021 (Figure 1). Potential sites were located using the iNaturalist (https://www.inaturalist.org/) and CalFlora (https://www.calflora.org/) databases. Seeds from 17 sites were collected (Appendix S2: Table S1). Seeds were collected from 27-30 individuals per site, located at least 1m apart along a linear transect, except for two sites with fewer collections (King Gillette Ranch, 16; Santa Cruz, 8). Seeds were stored at room temperature in paper envelopes until use.

**FIGURE 1.**
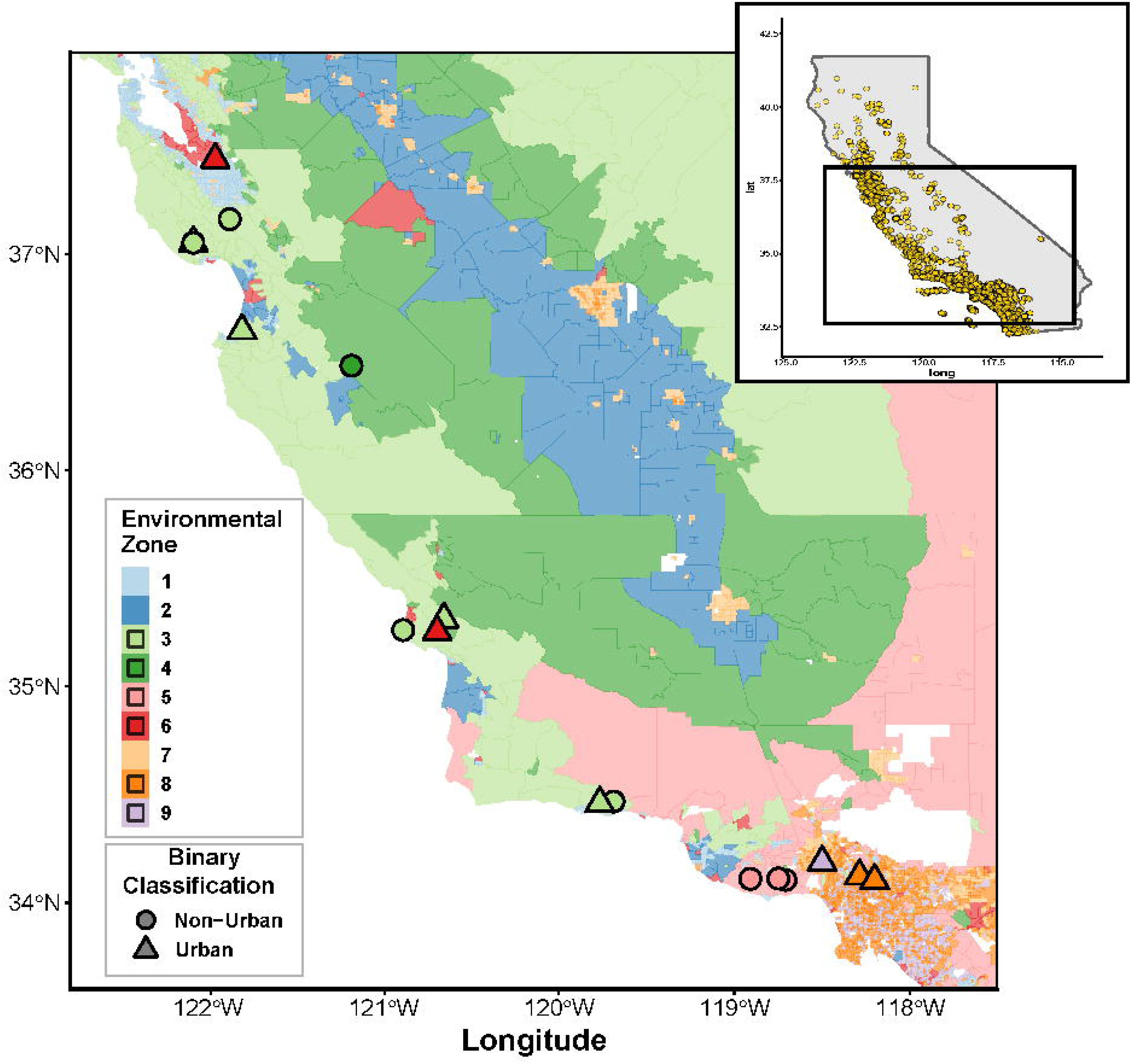
Map of California showing collection sites for Maltese starthistle, and their locations in six of the nine Environmental Zones (colors) identified in this study (see Figures 3 & 4). Collections were also independently identified as either Urban or Non-urban (symbols) based on the presence of human development at each site. Inset shows the current distribution of Maltese Starthisle (yellow dots) across the state of California obtained from the Global Biodiversity Information Facility (GBIF.org (24 June 2026) GBIF Occurrence Download https://doi.org/10.15468/dl.48njm5).

### Maltese starthistle trait measurements

To quantify trait differentiation of Maltese starthistle across Environmental Zones, resulting from evolved differences and/or plastic responses to seed environments, field-collected seeds were grown in a common greenhouse environment. Seeds from each maternal parent in the field were placed onto commercial potting soil (Happy Frog Potting Soil, FoxFarm Soil and Fertilizer Co, Arcata, CA, USA) under 12 h of fluorescent lights at 21 °C until they germinated. Individual plants were then transplanted into 410 ml Deepots (Steuwe & Sons Inc, Tangent, Oregon, USA) containers containing one of two different soil combinations: A) 1 part sand (30 grit Quikrete, The QUIKRETE Companies, Atlanta, GA, USA) to 5 parts soil (Happy Frog Potting Soil / coconut husk mixture (Chips-N-Fiber Premium Coconut Husk, PROCO, Tucson, Arizona, USA), hereafter the ‘soil’ treatment, or B) 1 part sand to 1 part soil / coconut husk mixture, hereafter the ‘sand’ treatment. Plants in the sand treatment were used for destructive root trait and specific leaf area measurements (see below). Plants were grown in the University of Arizona greenhouses (Tucson, AZ, USA) during the plant’s natural life cycle from October to April under ambient light and 24 °C average temperature. Final sample sizes for plants available for all measurements were 7 to 24 from each of the 17 sites in the soil treatment, and 7 to 16 from each of the 17 sites in the sand treatment.

We measured the length and width of the longest leaf, number of leaves, specific leaf area (SLA), root diameter and aboveground biomass. Reproductive traits measured were the number of flower heads (capitula) and the number of seeds produced by each flower head. Measuring SLA and root diameter is destructive, and these traits were measured in plants within the sand treatment. All other traits were measured on individuals in the soil treatment, which were allowed to grow undamaged through flowering. SLA was determined for a single leaf by measuring leaf areas with the Leafscan app (Anderson and Rosas-Anderson 2017. Leafscan Version 1.3.21. Retrieved from https://itunes.apple.com/app/id1254892230) followed by drying using silica gel for at least 3 days for dry mass measurements. The total number of flowers was recorded at the time of plant senescence. Finally, all aboveground biomass, excluding flower heads collected for seed collection, was collected and stored in brown paper bags. Samples were then oven-dried at 65 °C and weighed.

### Maltese starthistle trait differentiation across Environmental Zones

Each collection site was assigned an Environmental Zone from the cluster analysis, based on the census tract from which it was collected (Figure 1). Four sampling sites fell into census tracts that were excluded from the cluster analysis. These sites were manually assigned an Environmental Zone based on inspection of the Environmental Zones of census tracts around the site as well as the environmental characteristics known for the sampled site. We used linear models to test for trait differences among Environmental Zones, with Environmental Zone treated as a fixed categorical effect. Collection site was nested within Environmental Zone, as a nested fixed effect, to quantify any unique differences among sites. A separate linear model was used to predict each of the eight traits, with a Bonferroni correction of **α**/8 (= 0.006) applied when assessing the significance of each, to account for multiple tests of the same hypothesis of trait differentiation (Dunn 1961).

For comparison to existing approaches for identifying urban environments, we also conducted two additional sets of analyses of trait differentiation: i) A binary urban vs non-urban comparison that defines urban areas qualitatively, a traditional approach identified as pervasive in the literature review by Szulkin et al. (2020). We classified an environment as ‘Urban’ if it contained a large concentration of impervious surface or was visually among human structures and roadways (9 sites) and all other sites as ‘Non-urban’ (8 sites) (Figure 1; Appendix S2: Table S1), according to qualitative methods commonly found in urban eco-evolutionary literature (Murray-Stoker et al. 2025). ii) A three-category gradient based on percentage of impervious surface area (ISA) in the census tract containing each site. Census tracts including our sampling sites range from 0.2%-48.6% ISA (Appendix S2: Table S1). Our categories include tracts with <5% ISA that we identify is ‘Undeveloped’ (8 sites), as well as two ‘Developed’ categories based on those of the National Land Cover Database (NLCD) which designates <20% ISA as ‘Developed, Open Space’ (5 sites with 5-19% ISA), and 20-49% as ‘Developed, Low Intensity’ (4 sites). As for the analyses of traits by Environmental Zones, all analyses for these comparisons included fixed effects of category (whether binary or gradient) and nested fixed effects of collection site. Bonferroni corrections were again applied to assess significance.

## RESULTS

### Multivariate Environmental Zones

The PCA of spatial variables (Table 1) described a complex multivariate landscape of variation among census tracts, and reinforced that each of the variables provided largely unique information (Figure 2). PC Axes 1-4 explained 24.5%,12.5%, 8.2% and 7.7% of variation respectively, indicating that the dominant axes did not capture particularly large amounts of variation in the dataset and many dimensions were required to describe differences among census tracts. Multiple variables were correlated with the dominant axes (represented by long vectors), reflecting highly composite axes composed of several variables in order to explain variation. As expected by having filtered out variables with correlations above |0.7|, few variables were collinear in these dimensions. The strongest collinearities were between Impervious Surface and Diesel Particulate Matter in the PC1 vs PC2 space, and between Poverty Index and Asthma Rate in the PC3 vs PC4 space.

**FIGURE 2.**
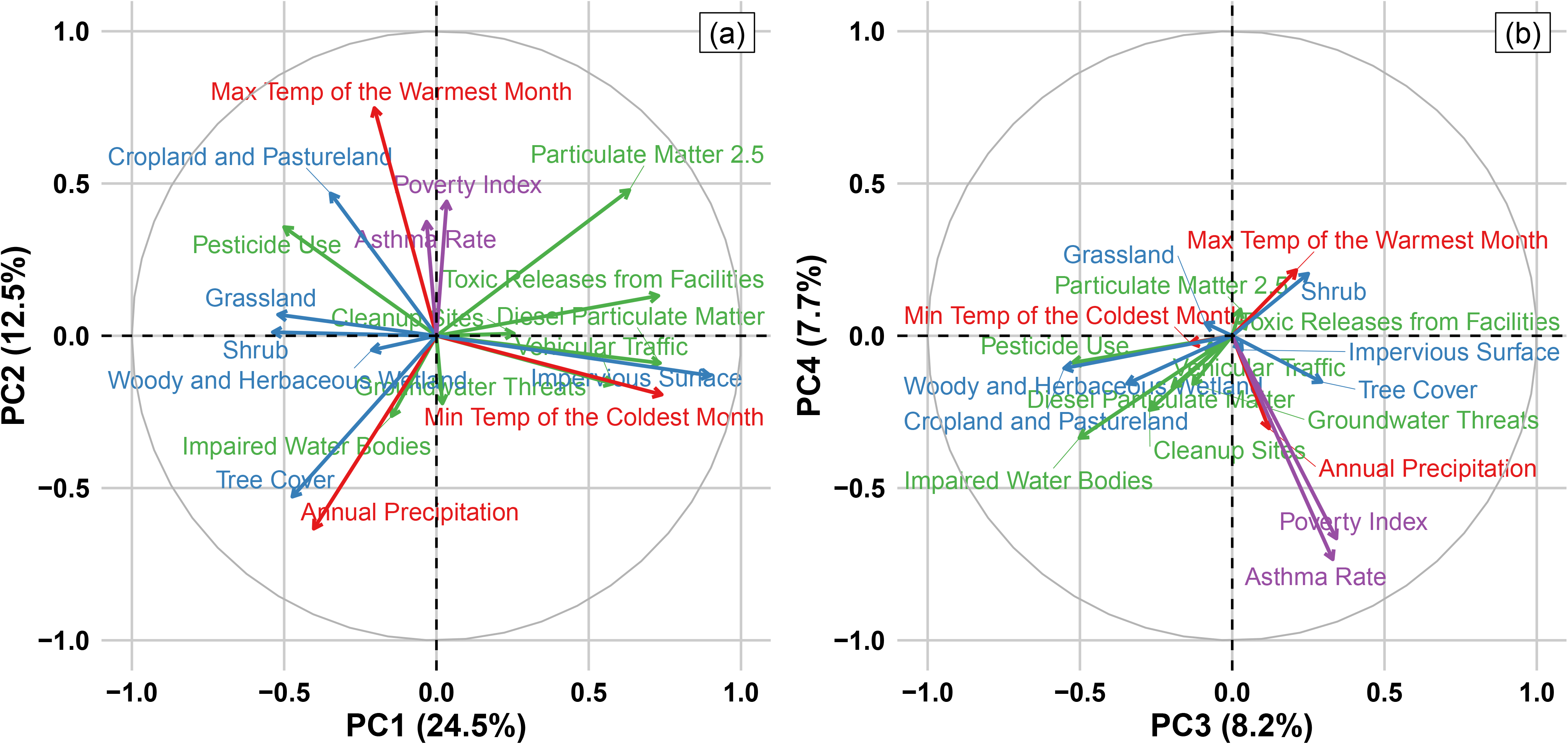
Principal Components Analysis of all 19 environmental variables used for the hierarchical cluster analysis, including climate (red), land cover (blue), pollution (green), and socio-economic (purple) variables. Shown are a) PC axes 1 and 2 and b) PC axes 3 and 4, with the percent of variation explained for each indicated in parentheses. Vectors show how variables (labeled) contribute to each axis (vector direction) and the strength of their correlation with these dimensions (vector length). Vectors reveal that most variables are contributing to these PC axes and each alone explains relatively little of the overall variation in this highly multivariate dataset.

Within the hierarchical cluster analysis, most clustering among census tracts occurred within the first 25% of distance in the dendrogram (i.e., most clustering occurred near the tips of the dendrogram, where distance between tract features was low), indicating that many tracts share similar multivariate features with groups of other tracts (Figure 3). Nine major clusters captured the majority of distance among branches in the dendrogram, reflecting deep divergences among different groups of closely clustered census tracts (Figure 3). These nine clusters, hereafter denoted Environmental Zones, were as follows: Environmental Zone 1: 1974 tracts (25.21%); Environmental Zone 2: 326 tracts (4.16%); Environmental Zone 3: 666 tracts (8.5%); Environmental Zone 4: 217 tracts (2.77%); Environmental Zone 5: 325 tracts (4.15%); Environmental Zone 6: 187 tracts (2.39%); Environmental Zone 7: 1595 tracts (20.37%); Environmental Zone 8: 1650 tracts (21.08%); Environmental Zone 9: 889 (11.36%).

**FIGURE 3.**
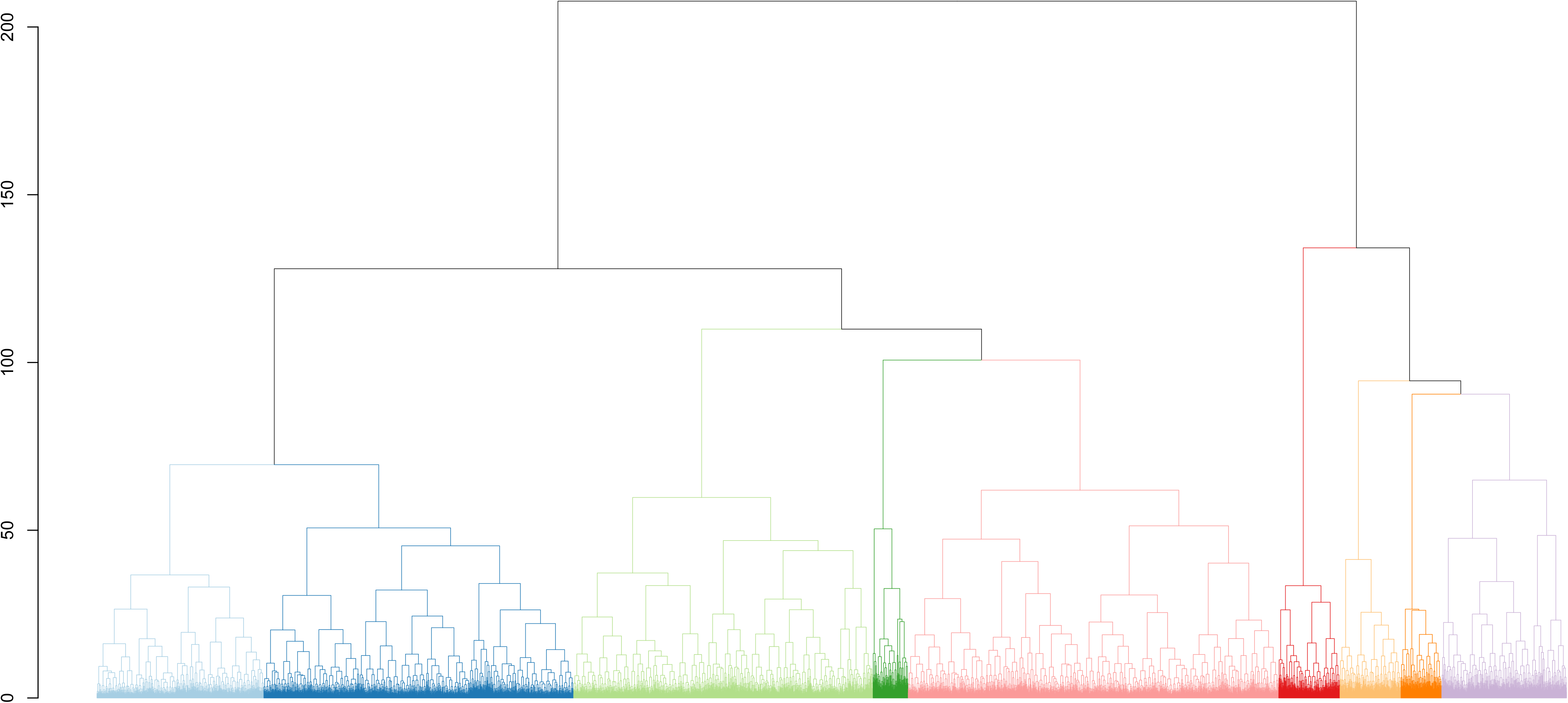
Hierarchical cluster analysis of 7,829 census tracts. Colors indicate membership in the nine Environmental Zones (numbered) that capture deep divergence among clusters. The y axis represents the Euclidean distance of branches between census tracts.

Environmental Zones revealed heterogeneity that deviated from common concepts of urban versus non-urban habitats. Figure 4 shows the most defining environmental variables for each of the nine Environmental Zones and a forest plot of the means of all 19 log-transformed variables used for the hierarchical clustering analysis, within each of the Environmental Zones that we identified, relative to their overall means. While Environmental Zones 1, 7, 8, and 9 contained the largest values for impervious surface, suggesting urban areas, Environmental Zone 1 was more closely clustered with Environmental Zones 2 - 5, which were associated with natural and agricultural land cover. Another common urbanization metric, tree cover, did not follow expected patterns in which high tree cover is associated with natural environments. For example, Environmental Zone 4 included natural areas with high levels of grassland and shrubland, but included overall less tree cover than Environmental Zone 1, 8, and 9 (all among the largest values for impervious surface). Additionally, Environmental Zones 3, 4, and 5 all contained high proportions of natural land cover types, but had differences in anthropogenic activity such as pesticide usage and waste clean up sites.

**FIGURE 4.**
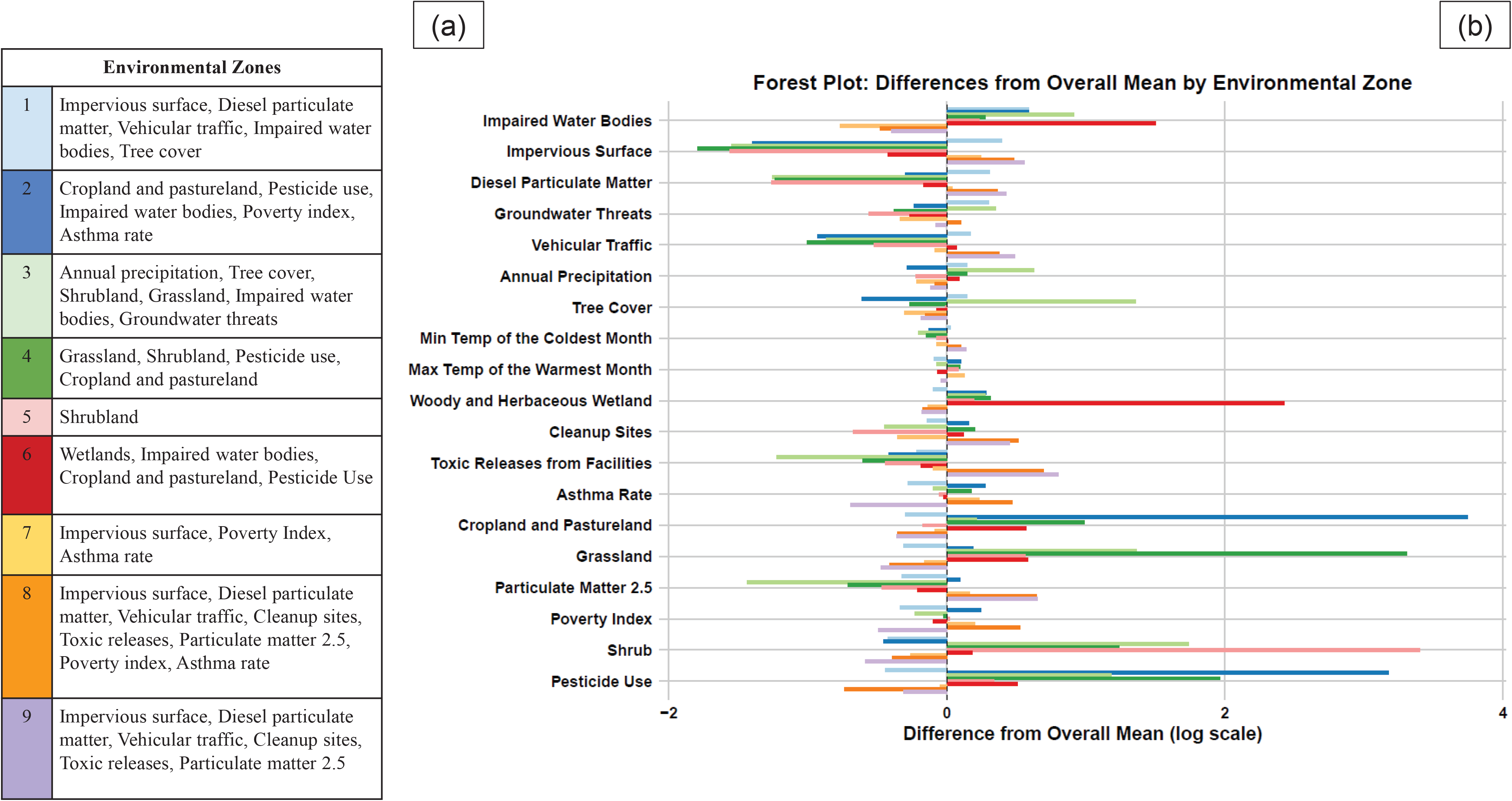
Forest plot of the mean values of all 19 environmental variables used in the hierarchical cluster analysis, for each of the nine Environmental Zones (colors). (a) Legend listing the environmental characteristics most associated with each of the nine Environmental Zones. (b) Forest plot showing differences in mean log-transformed values of the 19 environmental variables for each Environmental Zone. The x-axis represents the difference between the mean of log(x+1) for a variable within a specific Environmental Zone and the overall mean of log(x+1) across all 7,829 census tracts.

Areas containing major cities also differed in the abundance and distribution of different Environmental Zones (Figure 5; χ² = 3506.4, df = 16, p < 0.001). For example, Los Angeles County contained a greater proportion of Environmental Zone 8 than San Diego County and the San Francisco metropolitan areas combined. The “natural areas” surrounding urban cores in these areas differed qualitatively as well, with San Francisco’s urban centers being surrounded largely by a combination of Environmental Zones 3, 4, and 6, whereas the San Diego and Los Angeles outer areas were dominated by Environmental Zone 5 (Figure 5).

**FIGURE 5.**
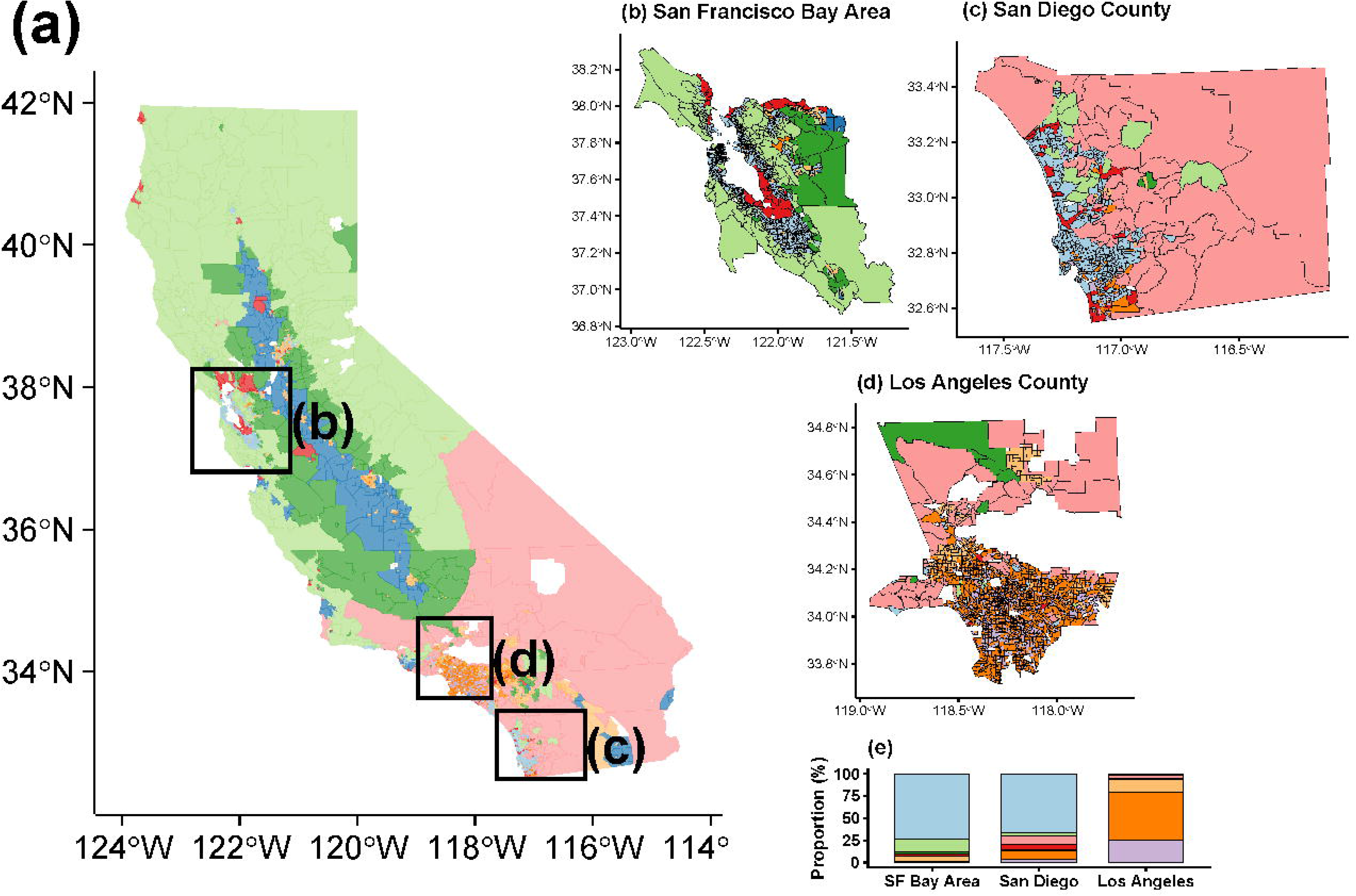
Environmental Zone classifications for census tracts of California. Polygons are state census tracts as of 2020, colored based on the Environmental Zones from the hierarchical cluster analysis (as described in Figures 3 & 4). Axes are latitude (x-axis) and longitude (y-axis). a) All 7,829 census tracts within the State of California included in this analysis. b) The metropolitan area of San Francisco Bay (1326 census tracts total). c) San Diego County (613 census tracts total). d) Los Angeles County (2251 census tracts total). e) The proportion of each Environmental Zone within each metropolitan area (χ² test: p<0.001)

The Environmental Zones also did not follow a pattern of urban-to-rural-to-natural gradients. Only Environmental Zone 3 showed a significant trend (increase) with distance to nearest urban center (Figure 6). For example, Environmental Zones 1, 7, 8, and 9 were all characterized by high anthropogenic activity (Figure 4), but only Zones 8 and 9 showed any evidence of decreasing as distance from the urban center increased, and this was not significant for either zone. Most zones more associated with natural environments also did not follow linear patterns. For instance, largely natural Environmental Zones 4 and 5 did not increase consistently as distance from urban centers increased; instead, they remained relatively constant at between 2-10% of land cover. Zone 3, associated with high tree cover and high precipitation, was the one exception, increasing significantly as distance to urban center increased. Finally, Environmental Zone 7 did not show a trend relative to urban centers, and tended to be highest at larger distances from them, despite containing qualities commonly associated with urban environments, such as relatively high Impervious Surface, Poverty Rate, and Asthma Index (Figure 4).

**FIGURE 6.**
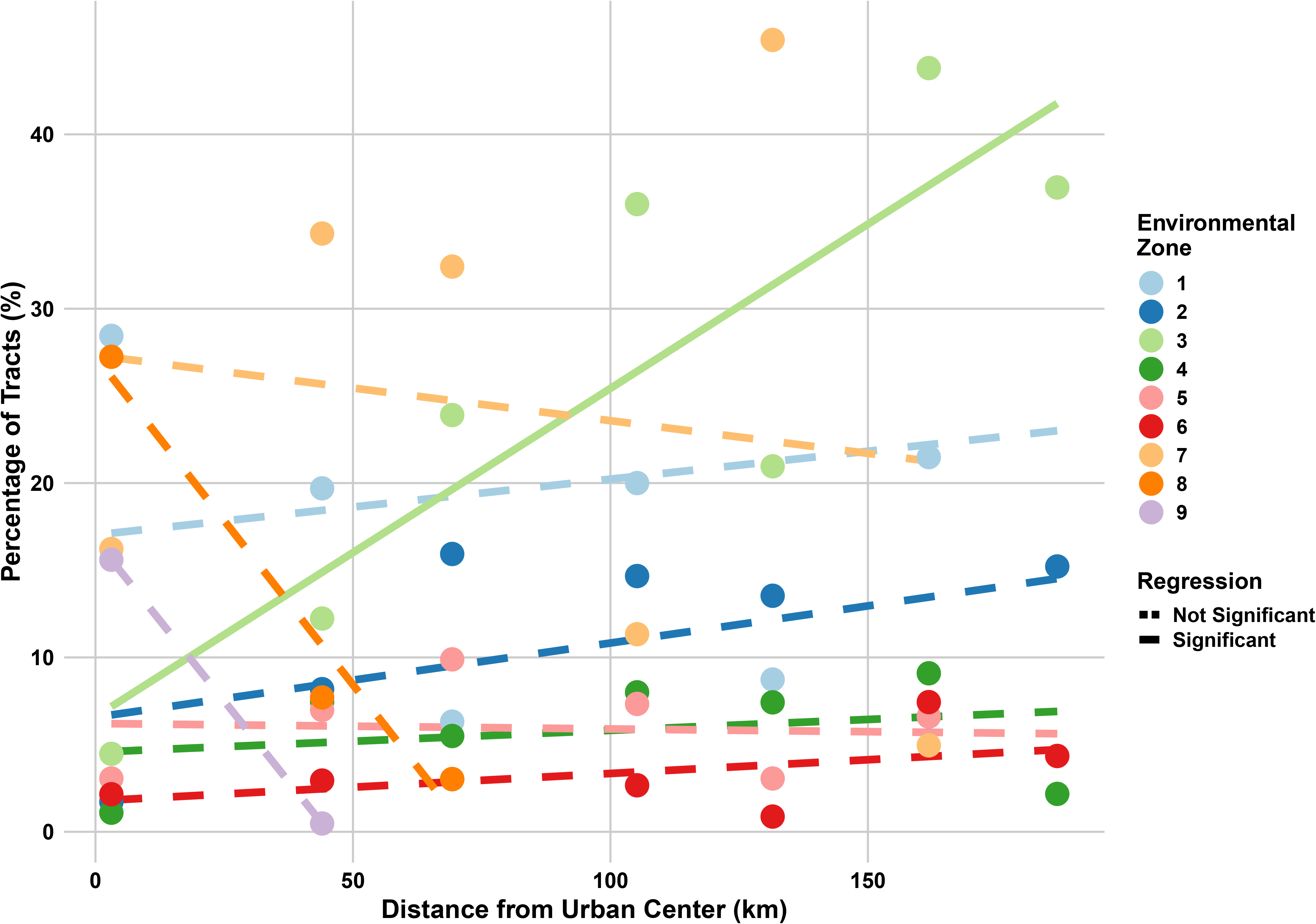
The percentage of tracts in each of the nine Environmental Zones (colors; as described in Figures 3 & 4) as a function of distance to the nearest urban center. Values on the x-axis represent the median distance of all census tracts within 30 km bins, with values on the y-axis indicating the percentage of tracts classified into each Environmental Zone within that bin. Lines represent linear regressions for each zone, with solid lines indicating statistically significant relationships (p < 0.05) and dashed lines indicating non-significant relationships. Distance was measured from the centroid of each census tract to the nearest continuous area of 80% or greater impervious surface that was greater than or equal to 2 km². Only Environmental Zone 3 showed a significant relationship (positive) with distance from urban centers (R² = 0.75, p = 0.011).

### Maltese starthistle trait differentiation

Maltese starthistle collection sites were assigned to six of the nine Environmental Zones (Figure 1), including Zones 3, 4, and 5 broadly associated with more natural land cover, Zones 8, and 9 broadly associated with more anthropogenic activities, and Zone 6 broadly associated with a mix of impacts (primarily wetlands and agriculture). Qualitative assessment during collection assigned the specific sites in Zones 4 and 5 as ‘Non-urban’ and sites in Zones 6, 8, and 9 as ‘Urban’ categories (Figure 1). Sites in Zone 3 were split between Urban and Non-Urban assignments (Figure 1). Each of the Urban sites within Zone 3 were associated with small developments adjacent to natural areas (Appendix S2: Table S1; pers. obs.), such that their census tract clustered with Zone 3 but traditional qualitative assignment identified them as urban.

When the binary urban land classification was used as the predictor, no Maltese starthistle traits differed significantly between Urban and Non-urban areas, before or after Bonferroni correction (Figure 7; Table 2; Appendix S2: Table S2). Nested effects of site were significant for six of the eight traits before correction, and for four after correction, indicating that at least half of traits did differ among sites (Appendix S2: Table S2). When the impervious surface gradient categories were used as a predictor, again, no Maltese starthistle traits differed significantly among categories before or after Bonferroni correction (Table 2; Appendix S2: Table S3; Appendix S2: Figure S1).

**FIGURE 7.**
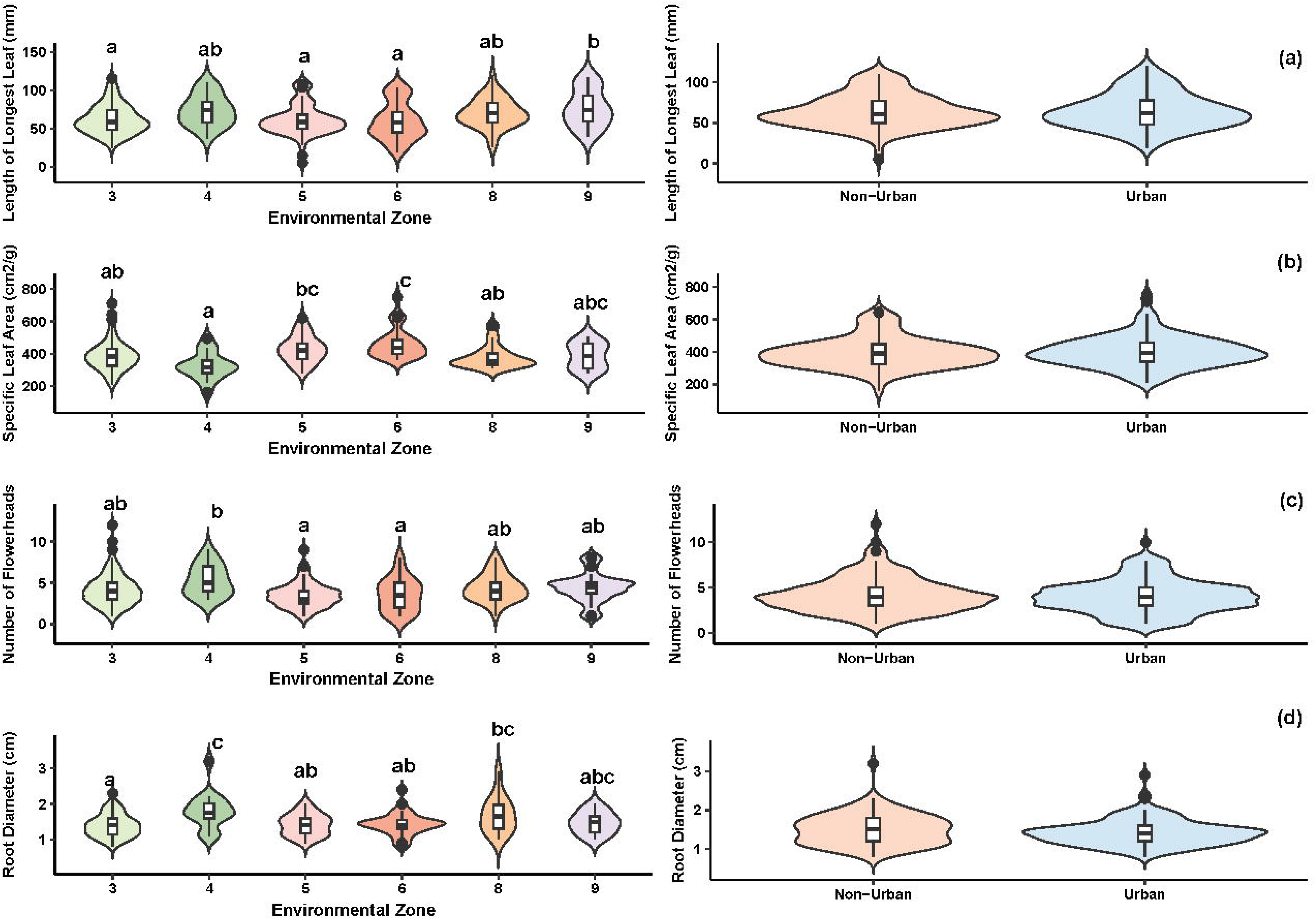
Tests for trait differentiation of Maltese starthistle in California for a) length of the longest leaf, b) specific leaf area (SLA), c) the number of flowering heads per plant, and d) root diameter. Shown are violin plots depicting the distribution of data, with boxplots inside indicating the median and quartiles. Letters above the violin plots indicate significantly different groups as determined by post hoc tests of linear models with main effects of either Environmental Zones (as described in Figures 3 & 4), or Urban vs Non-Urban environments. Seed collections occurred across six of the nine Environmental Zones identified across California.

**TABLE 2.** Linear models testing for trait differentiation of Maltese starthistle in California. Shown are *P*-values for the main effect in models predicting each trait, where effects were either the Environmental Zone assigned to each collection site, whether the site was Urban vs. Non-urban, or the Impervious Surface Gradient category the site was assigned. An asterisk denotes significant P-values after Bonferroni correction. Full model results are given in Supplementary Information Appendix 2 Tables S1 and S2.

| **Trait** | **Environmental**<br>**Zone**<br>**(*P*-value)** | **Urban vs. Non-Urban**<br>**(*P*-value)** | **Impervious Surface**<br>**Gradient**<br>**(*P*-value)** |
| --- | --- | --- | --- |
| Number of Leaves | 0.0717 | 0.5811 | 0.7135 |
| Width of Longest Leaf (mm) | 0.0095 | 0.0812 | 0.3365 |
| Length of Longest Leaf (mm) | 0.0006 * | 0.9398 | 0.7634 |
| Biomass (g) | 0.2420 | 0.0961 | 0.2395 |
| Root to Shoot Ratio | 0.0074 | 0.5913 | 0.4714 |
| Specific Leaf Area (cm<sup>2</sup>/g) | 1.8780e-06 * | 0.5692 | 0.0867 |
| Root Diameter (cm) | 9.4220e-05 * | 0.0948 | 0.0755 |
| Number of Flowerheads | 0.0024* | 0.2975 | 0.3002 |

In contrast, when Environmental Zones were used as the predictor, significant differences were observed in traits according to zones, including four after Bonferroni correction: length of longest leaf, SLA, root diameter, and the number of flowerheads (Figure 7; Table 2; Appendix S2: Table S4). Nested site effects within Environmental Types were statistically significant for three traits before correction, and only one (SLA) after (Appendix S2: Table S4). These analyses revealed differentiation between various types of natural / agricultural areas and more heavily urbanized areas, including thicker roots in plants from urban and impoverished Environmental Zone 8 relative to those from sites in the more natural Zone 3, and longer leaves in highly urban Zone 9 relative to less urban Zones 3/5/6. In addition, trait differences across more natural areas were also revealed, including plants from Zone 4 (dominated by grassland, shrubland, and agriculture) having lower SLA, larger root diameter, and more flowers than from other sites, and in contrast plants from Zone 6 (wetlands and agriculture) having higher SLA, smaller root diameter, fewer flowers, and shorter leaves.

## DISCUSSION

Our results highlight the importance of utilizing a multivariate classification system in urban ecological and evolutionary studies. The Environmental Zones that we identified within a single political unit captured a mosaic of environmental heterogeneity across census tracts within close geographic proximity and revealed new relationships between habitat type and the traits organisms exhibit. While the idea of the urban mosaic has been suggested by several authors (Ramalho & Hobbs, 2012; Szulkin et al., 2020; Winchell et al., 2022), few current land classifications have quantified them. Understanding this heterogeneity is increasingly recognized as essential to understanding both non adaptive and adaptive changes of organisms that inhabit urban spaces and beyond.

### Characterizing environmental heterogeneity in an urban mosaic

The nine Environmental Zones that we identified in California defined complex, nonlinear spatial distributions of urbanization, inconsistent with traditional dichotomies or univariate gradients for urban classification. For example, Environmental Zones 1, 7, 8 and 9 had the highest amounts of impervious surface area, but Zone 1 was more closely clustered (similar to) Zones 2-5, which contained more natural and agricultural environments, such as higher amounts of tree cover and grassland. This environmental heterogeneity was also evident in the principal component analysis of all variables, for which the dominant axes captured relatively little of the variation among census tracts, reflecting that many axes of multivariate variation were necessary to capture differences among tracts. This poses a challenge in that the resulting Environmental Zones cannot be well-described according to a small set of characteristics (a reason that we have avoided given the zones in our study names), but this underscores the reality that environmental variation can be fundamentally highly multivariate. Such extensive environmental heterogeneity across California makes it useful as a case study for methods to categorize heterogeneity, but we note that it may not reflect the degree of heterogeneity in other areas. Nevertheless, these results highlight the importance of avoiding simple classifications (such as extent of impervious surface) as indicators of urbanization when a mixture of other biologically important environmental factors are likely to be present, which is expected to be the case for most urbanized regions (Ramalho & Hobbs, 2012; Alberti et al., 2020; Miles et al., 2020; Murray-Stoker et al., 2025).

Furthermore, we observed that Environmental Zones were heavily interspersed across large metropolitan areas rather than having one or two zones dominating an “urban center” surrounded by increasingly natural environments. Although zones with the highest amount of development (8 and 9 in this study), were almost entirely located within 30 km of urban centers, zones containing high representation of impervious surface (e.g. Zones 1 and 7), varied in their distribution relative to the urban center, and no zone showed a significant decline as distance to an urban center increased. Additionally, the total percentage of census tracts labeled Zone 5, a zone associated with shrubland, remained relatively constant across distances from an urban center. Thus, the Environmental Zones in this study captured how natural areas can occur in relatively close proximity to cities, creating the urban mosaics recognized by previous studies calling for improvement in urban classifications (Pickett et al., 2016; Moll et al., 2019; Verrelli et al., 2022; Berg et al., 2024). The complex nature of these zones also illustrates that it is difficult to characterize

Another strength of the hierarchical cluster analysis used here is the opportunity to include important urban attributes such as socio-economic variables within classification models. The field of urban eco-evolutionary biology has been strongly urged to untangle the effects of socio-economics and socio-cultural variables on ecological and evolutionarily relevant traits in urban environments (Schell et al., 2020; Des Roches et al., 2021; Carlen et al., 2025). For instance, research on the luxury effect (the role of economic disparity in affecting urban flora composition) (Martin et al., 2004; Leong et al., 2018) and historic redlining (Schmidt & Garroway, 2022; Wood et al., 2024) have demonstrated that socio-economic inequality can impact eco-evolutionary mechanisms. However, while socio-economic variables have been associated with eco-evolutionary change (Peterson et al., 2020; Yitbarek et al., 2023; Estien et al., 2024), methods to integrate socio-economic data with other environmental data have been missing. Our study provides a demonstration of how these variables can be integrated with others. After excluding additional closely-correlated socio-economic variables, we included both Poverty Index and Asthma Rate in our clustering model and found important contributions of each to defining Environmental Zones. These results show how these variables can be used to define environmental classifications and support further study of the environmental differences associated with axes of socio-economic variation and their impact on urban flora and fauna.

### Comparing among cities

A major goal of urban ecology and evolutionary biology is to identify the common effects of urban environments across different cities. The idea of convergent evolution across urban environments has recently received extensive attention (Thompson et al., 2016; Diamond et al., 2018; Campbell-Staton et al., 2020). This idea predicts that traits of organisms inhabiting urban environments across multiple cities will converge in response to selection by a shared urban environment (Santangelo et al. 2020). However, tests of this hypothesis have had mixed results across cities and taxa (Yakub & Tiffin, 2017; Miles et al., 2025; Venegas et al., 2025). A lack of evidence for convergence could be due to substantial differences in the nature of urban environments across cities, such that areas defined as urban in each sample are not experiencing the same patterns of selection. While many cities do share key characteristics, including a high prevalence of impervious surface and human presence (Marzluff, 2001; Grimm et al., 2008; Johnson & Munshi-South, 2017), differences in environmental qualities such as climate, vegetation structure, land use, pollution, and socioeconomic variables may in turn alter selection on organisms within different cities (Alberti, 2023; Johnson et al., 2024; Carlen et al., 2025; Diamant et al., 2025). Thus, convergent evolution across cities may only be visible when it is possible to identify comparable types of environmental heterogeneity across cities, which may often be at smaller scales.

Our hierarchical cluster analysis highlights how even nearby urban centers can differ in their environmental variables and how environmental attributes within a single city can change over relatively small scales, making it difficult to compare similar environments among them. San Francisco, Los Angeles, and San Diego would all be considered dense urban metropolitan areas within coastal mediterranean-type climates, and they share similar histories of human colonization and development (Rolle & Verge, 2014; Schoenherr, 2017). Yet, these cities show clear differences in their distributions of the Environmental Zones that we identified. Los Angeles County was dominated by Environmental Zones 8 and 9, associated with higher particulate matter 2.5, toxic release from facilities, and clean-up sites. In contrast, San Diego and San Francisco were dominated by Environmental Zone 1 which had attributes such as a larger representation of impaired water bodies and higher rates of tree cover.

Comparisons among such cities that differ in their environmental composition will require a unified classification method that can define the urban mosaic in consistent ways across cities (Moll et al., 2019; Szulkin et al., 2020; Murray-Stoker et al., 2025). Our hierarchical cluster analysis offers an approach that can allow researchers to identify comparable environmental types within different urban areas, and also to make a priori predictions about which cities are most likely to exhibit convergent evolution of their biota, based on the similarity of biologically relevant environmental variables. With this information, researchers can plan sampling of comparable areas within and between urbanized regions, and test hypotheses about the specific environmental variables responsible for ecological and evolutionary change across cities.

### Trait variation in Maltese starthistle

Both within and across cities, the goal of urban ecology and evolution is often to understand how the urban environment affects the traits of organisms (Alberti et al., 2017; Diamond & Martin, 2021; Thompson et al., 2022). However, identifying the specific variables associated with trait variation has been difficult to disentangle within complex urban mosaics for the reasons outlined above (Rivkin et al., 2019; Verrelli et al., 2022; Winchell et al., 2023). Urbanization-related variables can vary independently of one another (Sanderfoot & Holloway, 2017; Grunst et al., 2020; Tüzün et al., 2025) and can impact environments at different spatial and temporal scales across the landscape (Uchida et al., 2021). Thus, a multivariate urban classification such as the hierarchical cluster analysis used here is needed to clarify associations between traits and multiple environmental variables.

As evidence of this, in our case study, trait differences in Maltese starthistle were captured using the Environmental Zones but not using the traditional dichotomous urban vs. non-urban qualitative classification, nor using a univariate gradient based on impervious surface area. Environmental Zones explained trait differences among the more heavily urbanized environments, with plants in Zone 9 (highest impervious surface) having longer leaves, while plants in Zone 8 (high impervious surface but greater poverty index) had thicker roots. One hypothesis for the differences seen in more impoverished areas is that these areas will be less irrigated in dry climates such as California, which may explain increased plant investment in roots in this zone (Hope et al., 2003; Leong et al., 2018; Diamant et al., 2025). The Environmental Zone classification also explained important differences in plant traits among more natural areas, with plants in drier zones having thicker leaves and roots, as well as more flowering heads, than plants in wetter zones. These trait differences are partially consistent with a ‘plant economics spectrum’ that predicts greater investment in roots and thicker leaves in resource-limited environments (Reich, 2014; de la Riva et al., 2021), though such limitations would not predict a greater number of flowering heads. Thus, our case study of Maltese starthistle trait variation raises several interesting new questions, which can be further investigated using experiments (such as reciprocal transplants or common gardens with different environmental treatments) to understand the fitness of different traits across these environmental zones (Lambert et al., 2021; Diamond et al., 2022; Fukano et al., 2023).

Half of the traits measured differed among Environmental Zones, with little variation among sites within zones, suggesting that Environmental Zones captured comparable environments affecting traits in different sites across California. An interesting illustration of how zones captured similarities across disparate sites occurred in Zone 3. We split sites in this zone into both ‘Urban’ and ‘Non-urban’ qualitative categories based on local observations of development in ‘Urban’ sites, but trait variation was better captured by grouping both types of sites into Zone 3. Our Zone 3 ‘Urban’ sites had close proximity to surrounding natural areas more typical of Zone 3, suggesting that for this species, traits were better characterized by the multivariate environment at the larger census tract scale, rather than the univariate level of development at the collection site. Thus, categorizing an appropriate scale of environmental zones prior to designing studies could inform researchers of optimal areas to sample for tests of specific *a priori* hypotheses, which has been identified as a need in urban ecology and evolutionary studies (Johnson & Munshi-South, 2017; Rivkin et al., 2019; Miles et al., 2020).

The presence or absence of an organism across zones could also be used to hypothesize patterns of habitat suitability. In this study, no populations of Maltese starthistle were collected in Environmental Zones 1, 2, and 7, associated with hyper-urbanization (particularly high vehicular activity) and agriculture. This could be due to either its absence from these areas or lack of collections. It raises the question of whether Maltese starthistle is unable to tolerate some of the most extreme aspects of urbanization, but may persist in other portions of cities and invade surrounding wildlands, a pattern theorized to explain the spread of invasive species that occupy urban environments (Francis & Chadwick, 2015; Padayachee et al., 2017; Borden & Flory, 2021).

## Conclusions

As the field of urban biology continues to grow (Johnson & Munshi-South, 2017; Szulkin et al., 2020; Collins et al., 2021) it is crucial that the variation in different aspects of urbanization is quantified (Ramalho & Hobbs, 2012; Miles et al., 2020; Schell et al., 2020; Winchell et al., 2022; Leandro et al., 2025). Without a standardized way to define urban areas and the heterogeneous environments within and around them, it is difficult to identify the impact of specific environmental variables on the ecology and evolution of organisms that inhabit anthropogenic spaces. Methods such as the hierarchical cluster analysis conducted here offer a way to identify multivariate environmental mosaics across the urban spectrum while also providing a framework for future sampling. Greater resolution of urban mosaics will allow for tests of predictions and hypotheses about biological variation across urbanized areas.

Our study demonstrates a powerful framework for testing biological questions across a heterogeneous landscape. It reveals opportunities to identify trait differences across complex urbanized landscapes that would not have been possible with univariate land classifications. Additionally, it demonstrates how environmental features can change over relatively short distances within urban spaces and vary at fine scales between them. This approach can thus benefit researchers interested in understanding the impact of heterogeneous environments on organisms inhabiting our increasingly urban world.

## ACKNOWLEDGMENTS

We would like to thank J. Autz, R. Rives, M. Carr, S. Lockhart, and C. Chance for assistance in the greenhouse. In addition, we would like to thank the Urban Research Center from the Natural History Museum of Los Angeles County for their advice on project design. Finally, we thank J. Bronstein and two anonymous reviewers for helpful comments on earlier versions of this manuscript.

## AUTHOR CONTRIBUTIONS

Anthony Dant conceived the study. Anthony Dant and Katrina Dlugosch designed and secured funding for the research. Anthony Dant and Lily Bishop conducted the research. Anthony Dant conducted analyses and drafted the manuscript, with assistance from Katrina Dlugosch. All authors reviewed and approved the final version of the manuscript

