## Appendix 1 for "A classification approach to categorize urbanized landscapes in California and trait variation in Maltese starthistle"

**Appendix S1 for:**

Title: A classification approach to categorize urbanized landscapes in California and trait variation in Maltese starthistle

Authors: Anthony J. Dant, Lily G. Bishop, Katrina M. Dlugosch

Journal: Ecological Applications

**Supporting Methods - Spatial variable selection**
Spatial variables of four types were considered: Climatic, Pollution, Socio-economic, and Land Cover. For each of these, available variables were considered and either retained or excluded as follows:

<u>1. Climatic Variables</u>
Source for all climate variables: WorldClim Database
19 bioclim variables available, 3 retained

Excluded:
- 16 variables excluded: All other variables had high correlations with the three selected variables (see Figures S1 and S2 below).

Retained:
- Annual Mean Temperature
- Maximum Temperature of Warmest Month
- Annual Mean Precipitation

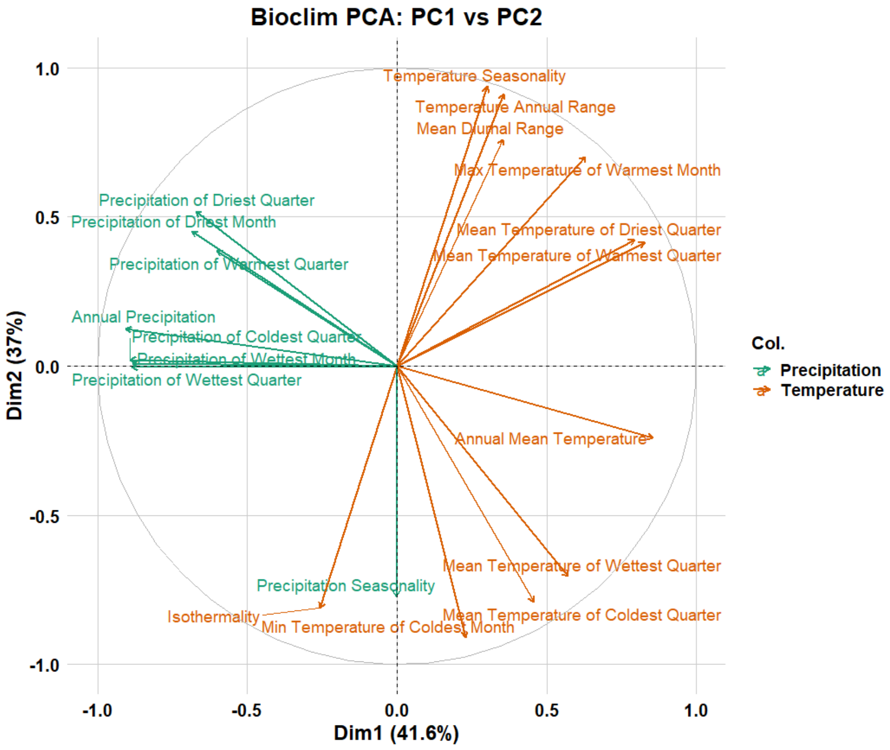


**Fig S1.** Principle Components Analysis of all 19 BioClim variables for each of the census tracts in California. Colinearities among many variables are evident.

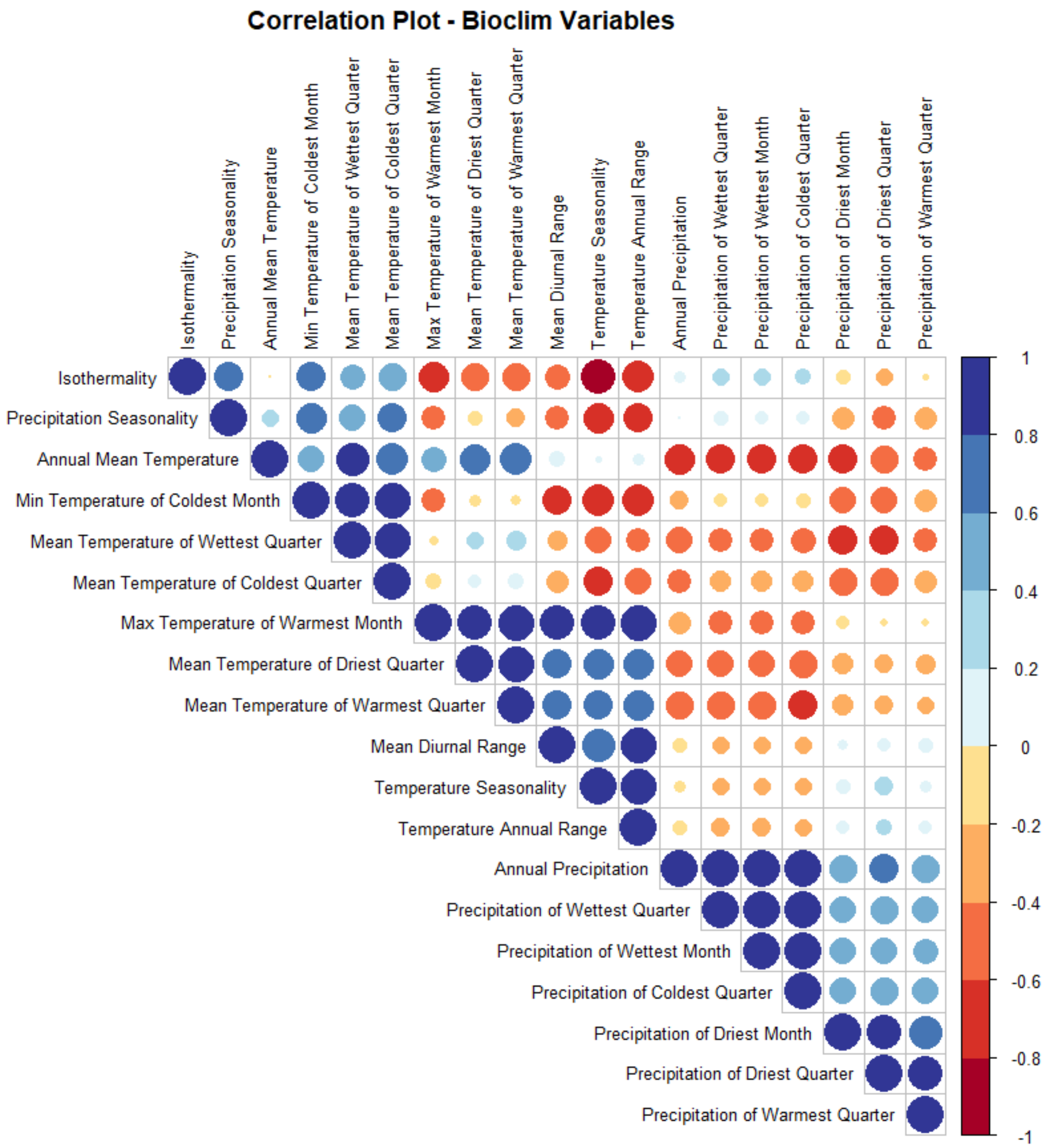


**Fig S2.** Correlation analysis of all 19 BioClim variables for each of the census tracts in California. Correlations among many variables are evident, as is the independence (correlations near zero) among the three variables retained.

<u>2. Pollution Variables</u>
Source for all pollution variables: 2019 CalEnviroScreen 4.0 Report (CALEPA)
13 variables available, 8 retained

Excluded due to low relevance for wild organisms:

- Lead Percentile: Potential risk for lead exposure in children living in low-income communities with older housing.
- Drinking Water Percentile: Drinking water contaminant index for selected contaminants (2011 to 2019).
- HazWasteP: Sum of weighted permitted hazardous waste facilities, hazardous waste generators, and chrome plating facilities within each census tract.
- SolWasteP: Sum of weighted solid waste sites and facilities (as of July 2021).

Excluded due to correlations with other variables:

- Ozone Percentile: Mean of summer months (May-October) of the daily maximum 8-hour ozone concentration (ppm), averaged over three years (2017 to 2019). Reason for removal: highly correlated with Max Temperature of Warmest Month (over 0.7%).

Retained:

- Particulate matter 2.5: Annual mean concentration of PM2.5 (weighted average of measured monitor concentrations and satellite observations, µg/m3), over three years (2015 to 2017).
- Pesticide usage: Specific pesticides included in the measure of pesticide use were narrowed from the list of all registered pesticides in use in California to focus on a subset of 132 chemicals that are filtered for hazard and volatility.
- Vehicular traffic: Traffic impacts represent the vehicles in a specified area, resulting in human exposures to chemicals that are released into the air by vehicle exhaust, as well as other effects related to large concentrations of motor vehicles.
- CleanupP: Sites undergoing cleanup actions by governmental authorities or by property owners that have suffered environmental degradation due to the presence of hazardous substances.
- Ground Water Threat Percentile: Common groundwater pollutants found at LUST and cleanup sites in California including gasoline and diesel fuels, chlorinated solvents and other volatile organic compounds (VOCs).
- Impaired Water Bodies Percentile: Summed number of pollutants across all water bodies designated as impaired within the area (2018).
- Toxic_Rel_P: Toxic Release Inventory (TRI) of on-site releases to air, water, and land and underground injection of any classified chemical, as well as quantities transferred off-site.
- DieselPM_P: Gridded diesel PM emissions from on-road sources.

<u>3. Socio Economic Variables</u>
Source for all socio-economic variables: 2019 CalEnviroScreen 4.0 Report (CALEPA)
7 variables available, 2 retained

Excluded due to missing data; inclusion would mean the removal of a large portion of census tracts:

- Unemployment Percentile: Percentage of the population over the age of 16 that is unemployed and eligible for the labor force.
- Linguistic Isolation Percentile: Percentage of limited English-speaking households, (2015-2019).
- Housing Burden Percentile: Percent of households in a census tract that are both low income (making less than 80% of the HUD Area Median Family Income) and severely burdened by housing costs (paying greater than 50% of their income to housing costs).
- Educational attainment Percentile: Percentage of the population over age 25 with less than a high school education (5-year estimate, 2015-2019).

Excluded due to redundancy with census tract delineation:

- Total Population: From 2020 Census

Retained:

- Poverty Rate Percentile: Percent of the population living below two times the federal poverty level (5-year estimate, 2015-2019)
- Asthma Rate Percentile: Spatially modeled, age-adjusted rate of emergency department visits for asthma per 10,000 (averaged over 2015-2017).

4. Land Cover Variables

Source for all land cover variables: United States National Land Cover Database, including:

- Annual Land Cover database: U.S. Geological Survey (USGS), 2024, Annual NLCD Collection 1 Science Products: U.S. Geological Survey data release, https://doi.org/10.5066/P94UXNTS.
- Treecover: Survey, U.G., and Service, U.F., 2024, *National Land Cover Database (NLCD) Tree Canopy Cover Products*: U.S. Geological Survey data release, https://doi.org/10.5066/P9JZ7AO3.
- Impervious Surface: U.S. Geological Survey (USGS), 2024, Annual NLCD Collection 1 Science Products: U.S. Geological Survey data release, https://doi.org/10.5066/P94UXNTS
- Available variables include those in the table shown below. Definitions for each variable (obtained from the legends of NLCD) are shown below the table

Excluded:

- 'Water' was excluded as a variable, so that tracts would be clustered based on their terrestrial composition.
- 'Developed' land cover was replaced with % of Total Impervious Surface from the Annual NLCD Collection 1 Science Products: U.S. Geological Survey
- Forest was replaced with % Tree Cover from the National Land Cover Database (NLCD) Tree Canopy Cover Products
- 'Barren' land was also excluded from this analysis due to lack of habitat, such that tracts would be characterized by other available habitat and/or impervious surface in the area.

Retained:

- Tree Cover: Total Tree Cover as a Percentage (0-100%)
- Impervious Surface: Total Impervious Surface as a Percentage (0-100%)
- Grassland: Grassland/Herbaceous- areas dominated by gramanoid or herbaceous vegetation, generally greater than 80% of total vegetation. These areas are not subject to intensive management such as tilling, but can be utilized for grazing.
- Shrubland: areas dominated by shrubs; less than 5 meters tall with shrub canopy typically greater than 20% of total vegetation. This class includes true shrubs, young trees in an early successional stage or trees stunted from environmental conditions.
- Planted/Cultivated: Combined the following:
    - Pasture/Hay: Areas of grasses, legumes, or grass-legume mixtures planted for livestock grazing or the production of seed or hay crops, typically on a perennial cycle. Pasture/hay vegetation accounts for greater than 20% of total vegetation.
    - Cultivated Crops: Areas used for the production of annual crops, such as corn, soybeans, vegetables, tobacco, and cotton, and also perennial woody crops such as orchards and vineyards. Crop vegetation accounts for greater than 20% of total vegetation. This class also includes all land being actively tilled.
- Wetlands_Percent: Combined the following:

- Woody Wetlands: Areas where forest or shrubland vegetation accounts for greater than 20% of vegetative cover and the soil or substrate is periodically saturated with or covered with water.
- Emergent Herbaceous Wetlands: Areas where perennial herbaceous vegetation accounts for greater than 80% of vegetative cover and the soil or substrate is periodically saturated with or covered with water.
