## Appendix 2 for "A classification approach to categorize urbanized landscapes in California and trait variation in Maltese starthistle"

**Appendix S2 for:**

Title: A classification approach to categorize urbanized landscapes in California and trait variation in Maltese starthistle

Authors: Anthony J. Dant, Lily G. Bishop, Katrina M. Dlugosch

Journal: Ecological Applications

**Supporting Methods - Distance to urban center**

The distance of census tracts to their nearest urban center was calculated in the following way using [ArcGIS Pro/ version 3.5.0]:

1. NLCD raster was converted to binary (1 = class 24, 0 = all else) to isolate high-intensity urban pixels using ‘Reclassify NLCD’.
2. Binary raster was converted to polygons for area-based filtering using ‘Raster to Polygon’.
3. Polygon area was calculated in square kilometers using ‘Calculate Area’ and saved as ‘Area_km2’.
4. Polygons were filtered to keep only those with Area_km2 greater than or equal to 2.0 using ‘Select Urban Centers’.
5. Selected polygons were exported as 'Urban_Centers_2km2' using ‘Export Selection’.
6. Distance was calculated from 'Urban_Cores_2km2' with max distance 200,000 meters and 30m cell size using ‘Euclidean Distance’.
7. Minimum distance of each tract was extracted using census tracts as zone features and distance raster as value raster in ‘Zonal Statistics as Table’.
8. Each tract was associated with its nearest urban core distance by joining Zonal Statistics output back to tract shapefile using ‘Join Table’.

**Table S1.** *Centaurea melitensis* collection sites in California. The '# of Individuals Collected' refers to the number of maternal individuals from whom seed was collected along a transect at each site. Binary Classifications are based on whether collections sites were located in areas of human development (Urban) or in undeveloped areas (Non-urban). Impervious Surface Percentage is calculated for the census tract in which the collection site occurred.

| **Location Name** | **Nearest City** | **# of Individuals Collected** | **Latitude Longitude** | **Binary Classification** | **Impervious Surface Percentage** |
|---|---|---|---|---|---|
| Reagan Ranch | Los Angeles | 30 | 34°06'30"N 118°44'57"W | Non-Urban | 2.1% |
| Lake Balboa | Los Angeles | 30 | 34°11'01"N 118°29'59"W | Urban | 24.4% |
| Ernest Debbs Park | Los Angeles | 30 | 34°05'58"N 118°12'03"W | Urban | 26.1% |
| Griffith Park | Los Angeles | 30 | 34°07'13"N 118°17'13"W | Urban | 13.8% |
| King Gillette Ranch | Los Angeles | 16 | 34°06'08"N 118°42'33"W | Non-Urban | 7.5% |
| Sandstone Peak | Los Angeles | 30 | 34°06'26"N 118°54'33"W | Non-Urban | 1.2% |
| San Marcos Park | Santa Barbra | 30 | 34°27'24"N 119°46'00"W | Urban | 2.8% |
| Rattlesnake Canyon | Santa Barbra | 30 | 34°27'53"N 119°41'17"W | Non-Urban | 4.1% |
| Cal Poly San Luis Obispo | San Luis Obispo | 30 | 35°18'32"N 120°39'17"W | Urban | 6.3% |
| Irish foothills | San Luis Obispo | 30 | 35°15'05"N 120°41'36"W | Urban | 20.3% |
| Montana de Oro State Park | San Luis Obispo | 27 | 35°15'29"N 120°53'14"W | Non-Urban | 0.3% |
| Pinnacles National Park | Monterey | 30 | 36°28'53"N 121°10'54"W | Non-Urban | 0.2% |
| Seaside, Monterey | Monterey | 30 | 36°38'38"N 121°48'19"W | Urban | 48.6% |

| | | | | | |
|---|---|---|---|---|---|
| UC Santa Cruz | Santa Cruz | 30 | 36°59'55"N 122°03'23"W | Urban | 11.4% |
| Henry Cowell, Redwood State Park | Santa Cruz | 8 | 37°02'59"N 122°04'59"W | Non-Urban | 1.7% |
| Sierra Azul Open Space Reserve | Santa Cruz | 30 | 37°09'34"N 121°52'33"W | Non-Urban | 0.9% |
| Don Edwards Reserve | San Jose | 30 | 37°26'08"N 121°57'29"W | Urban | 14.7% |

**Table S2.** Linear models testing for trait differentiation of *Centaurea melitensis* in California, with Urban vs Non-Urban Land Classification as the main effect, and a nested site effect. An asterix denotes significant *P*-values after Bonferroni correction.

| Trait | Effect | df | df residual | F | *P* |
|---|---|---|---|---|---|
| Number of Leaves | Urban vs. Non-Urban | 1 | 292 | 0.3052 | 0.58107 |
| | Site (nested) | 15 | 292 | 2.1606 | 0.00774 |
| Width of Longest Leaf (mm) | Urban vs. Non-Urban | 1 | 292 | 3.0629 | 0.08115 |
| | Site (nested) | 15 | 292 | 1.4697 | 0.11533 |
| Length of Longest Leaf (mm) | Urban vs. Non-Urban | 1 | 292 | 0.0057 | 0.93983 |
| | Site (nested) | 15 | 292 | 2.7945 | 0.00045* |
| Biomass (g) | Urban vs. Non-Urban | 1 | 280 | 2.7887 | 0.09605 |
| | Site (nested) | 15 | 280 | 1.3880 | 0.15206 |
| Root to Shoot Ratio | Urban vs. Non-Urban | 1 | 188 | 0.2893 | 0.59129 |
| | Site (nested) | 15 | 188 | 1.9637 | 0.01982 |
| Specific Leaf Area ($cm^2/g$) | Urban vs. Non-Urban | 1 | 194 | 0.3252 | 0.56920 |
| | Site (nested) | 15 | 194 | 5.7966 | 8.035e-10* |
| Root Diameter (cm) | Urban vs. Non-Urban | 1 | 199 | 2.8175 | 0.094810 |
| | Site (nested) | 15 | 199 | 2.4226 | 0.00295* |
| Number of Flowerheads | Urban vs. Non-Urban | 1 | 289 | 1.0892 | 0.29752 |
| | Site (nested) | 15 | 289 | 2.4911 | 0.00182* |

**Table S3.** Linear models testing for trait differentiation of *Centaurea melitensis* in California, with Impervious Surface Gradient category as the main effect, and a nested site effect. An asterix denotes significant *P*-values after Bonferroni correction.

| Trait | Effect | df | df residual | F | *P* |
|---|---|---|---|---|---|
| Number of Leaves | Impervious Surface% | 2 | 292 | 0.3379 | 0.71358 |
| | Site (nested) | 14 | 292 | 2.2884 | 0.00552* |
| Width of Longest Leaf (mm) | Impervious Surface% | 2 | 292 | 1.0932 | 0.3365 |
| | Site (nested) | 14 | 292 | 1.6373 | 0.0686 |
| Length of Longest Leaf (mm) | Impervious Surface% | 2 | 292 | 0.2702 | 0.76342 |
| | Site (nested) | 14 | 292 | 2.9559 | 0.00031* |
| Biomass (g) | Impervious Surface% | 2 | 280 | 1.4364 | 0.23952 |
| | Site (nested) | 14 | 280 | 1.4811 | 0.11721 |
| Root to Shoot Ratio | Impervious Surface% | 2 | 188 | 0.7549 | 0.47146 |
| | Site (nested) | 14 | 188 | 2.0168 | 0.01852 |
| Specific Leaf Area (cm^2/g) | Impervious Surface% | 2 | 194 | 2.47683 | 0.08666 |
| | Site (nested) | 14 | 194 | 5.8801 | <0.00001* |
| Root Diameter (cm) | Impervious Surface% | 2 | 199 | 2.6180 | 0.07546 |

| | | | | | |
|---|---|---|---|---|---|
| | Site (nested) | 14 | 199 | 2.4228 | 0.0037* |
| Number of Flowerheads | Impervious Surface% | 2 | 289 | 1.2082 | 0.30023 |
| | Site (nested) | 14 | 289 | 2.57419 | 0.00165* |

**Table S4.** Linear models testing for trait differentiation of *Centaurea melitensis* in California, with Environmental Zone as the main effect, and a nested site effect. An asterix denotes significant *P*-values after Bonferroni correction.

| Trait | Effect | df | df residual | F | *P* |
|---|---|---|---|---|---|
| Number of Leaves | Environmental Zone | 5 | 292 | 2.0366 | 0.07354 |
| | Site (nested) | 11 | 292 | 2.0483 | 0.02410 |
| Width of Longest Leaf (mm) | Environmental Zone | 5 | 292 | 3.1086 | 0.00946 |
| | Site (nested) | 11 | 292 | 0.8696 | 0.57067 |
| Length of Longest Leaf (mm) | Environmental Zone | 5 | 292 | 4.4820 | 0.00060* |
| | Site (nested) | 11 | 292 | 1.7739 | 0.05799 |
| Biomass (g) | Environmental Zone | 5 | 280 | 1.3335 | 0.25010 |
| | Site (nested) | 11 | 280 | 1.5401 | 0.11670 |
| Root to Shoot Ratio | Environmental Zone | 5 | 188 | 3.2694 | 0.00744 |
| | Site (nested) | 11 | 188 | 1.2180 | 0.27743 |
| Specific Leaf Area (cm^2/g) | Environmental Zone | 5 | 194 | 7.4935 | 1.878e-06* |

| | | | | | |
|---|---|---|---|---|---|
| | Site (nested) | 11 | 194 | 4.5279 | 4.235e-06* |
| Root Diameter (cm) | Environmental Zone | 5 | 199 | 5.4838 | 9.422e-05* |
| | Site (nested) | 11 | 199 | 1.0670 | 0.38980 |
| Number of Flowerheads | Environmental Zone | 5 | 289 | 3.7877 | 0.00244* |
| | Site (nested) | 11 | 289 | 1.7742 | 0.05798 |

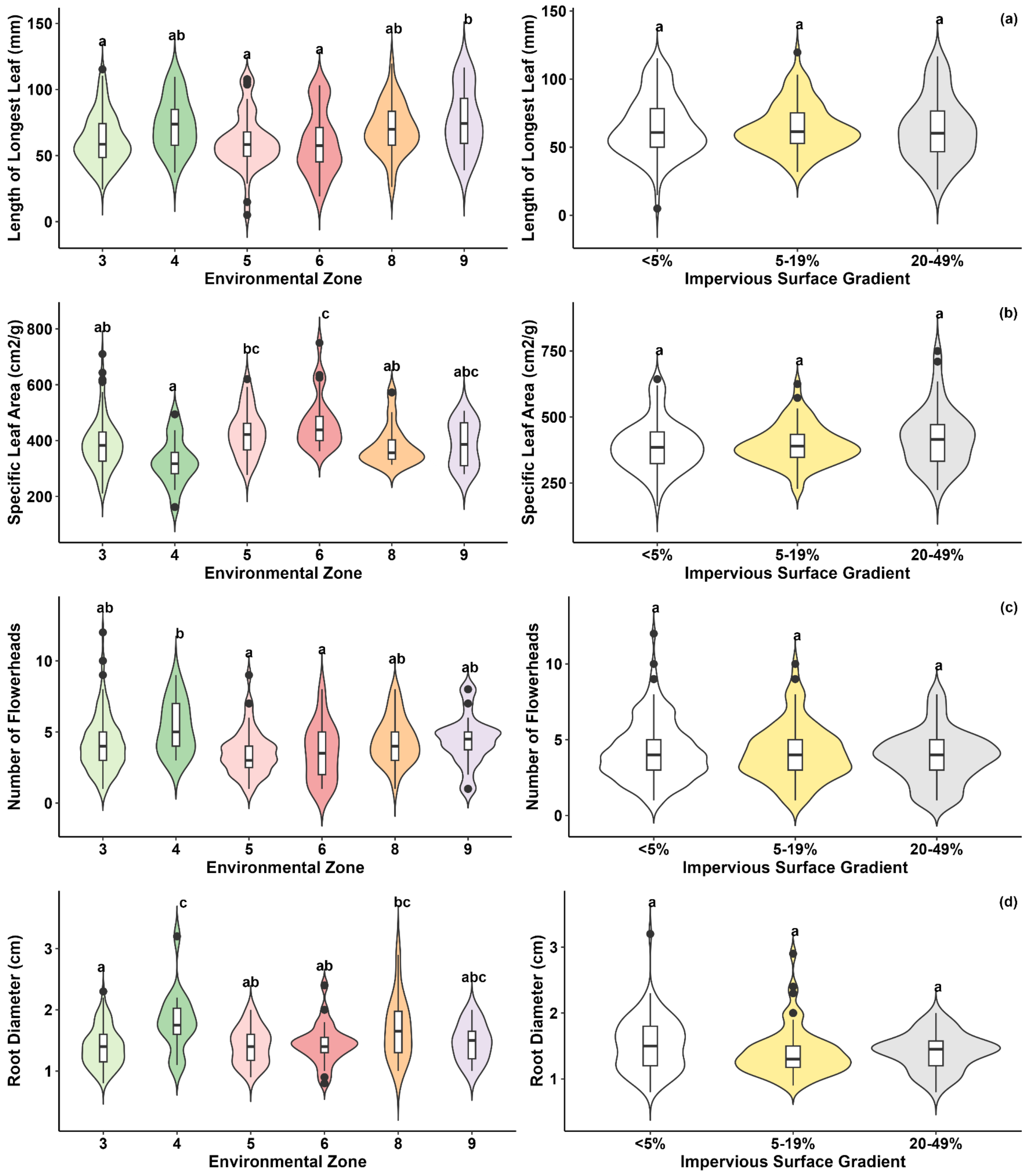


**Figure S1.** Tests for trait differentiation of Maltese starthistle in California for a) length of the longest leaf, b) specific leaf area (SLA), c) the number of flowering heads per plant, and d) root diameter. Shown are violin plots depicting the distribution of data, with boxplots inside indicating the median and quartiles. Letters above the violin plots indicate significantly different groups as determined by post hoc tests of linear models with main effects of either Environmental Zones, or Impervious Surface Gradient categories.
